# Koisio Technology-Modulated Cell Culture Media can Lead to Markedly Increased Intracellular ATP Levels of Fibroblast Cell Cultures

**DOI:** 10.1101/2022.02.28.482249

**Authors:** Yinghui Men, Mingchao Zhang, Qi Zhu, Weihai Ying

## Abstract

It has been reported that Koisio technology-modulated water without external additions of any substances can enter several types of aquaporins with significantly higher rate. In this study we determined the effects of Koisio technology-modulated cell culture media (Koisio media) on the energy state of cells. We found that for the lung fibroblast (L929) cell cultures, Koisio media without any additions of external substances led to approximately an 57% increase in the intracellular ATP levels of the cell cultures at the Cell Passage 6 - 18, compared with regular cell culture media. Seemingly surprisingly, we did not find that Koisio media could significantly affect the mitochondrial membrane potential of the cells, compared with regular cell culture media. Collectively, our study has provided first evidence indicating that Koisio media can lead to remarkable increases in the intracellular ATP levels of the L929 cell cultures, suggesting significant potential of Koisio technology in generating energy-enhancing water. Our finding that Koisio media did not affect the mitochondrial membrane potential has suggested that novel mechanisms may underlie the effects of Koisio media on cellular energy.

## Introduction

Intracellular ATP plays crucial roles in numerous cellular functions (Stryer, 1995). Therefore, maintenance of intracellular ATP levels is critical for cellular functions and cell survival (Almeida and Bolanos, 2001, Beal, 1992, Eguchi et al., 1997, Feldenberg et al., 1999, Stryer, 1995). Intracellular ATP in normal cells is generated mostly through mitochondrial oxidative phosphorylation, with a small portion of intracellular ATP being generated through glycolysis (Stryer, 1995). Intracellular ATP levels are regulated by numerous factors, failures of which can lead to a number of pathological changes (Almeida and Bolanos, 2001, Beal, 1992, Eguchi et al., 1997, Feldenberg et al., 1999, Stryer, 1995, Williams et al., 2015). Together with oxidative stress and inflammation, defects of energy metabolism belong to one of the most critical pathological factors of aging and numerous diseases (Beal, 1995, Ying, 1996a, 1996b, 1997a, 1997b).

It is highly valuable to discover novel and economic substances for enhancing the energy levels of human body. Since water drinking is required for human survival, it would be of great value if we can produce certain type of water with capacity of enhancing the energy levels of human body without addition of external substances. However, it is exceedingly difficult to reach this goal.

A previous study has reported that the ceramics produced by Koisio technology could produce water with increased permeability through aquaporins (Kozumi and Kitagawa, 2016). However, it has been unclear if Koisio technology-modulated solutions without additions of any external substances may affect intracellular ATP levels. In current study, we determined the effect of Koisio technology-modulated solutions on intracellular ATP levels, showing that Koisio media can produce remarkable increases in the intracellular ATP levels of fibroblast cell cultures.

## Materials and Methods

### Cell Cultures

L929 cells were purchased from the national collection of authenticated cell cultures (Shanghai, China). Cells were incubated in Dulbecco’s modified Eagle medium (HyClone, Logan, UT, United States) supplemented with 10% fetal bovine serum (Gibco, Carlsbad, CA, United States), 1% 100 U/ml penicillin, and 100 μg/ml streptomycin at 37 °C in a humidified incubator under 5% CO_2_.

### Modulations of cell culture media by Koisio technology

Modulations of cell culture media were conducted by Koisio technology as reported previously (Kozumi and Kitagawa, 2016).

### Determinations of intracellular ATP levels

As described previously (Zhang et al., 2018), cells in 24-well plates were harvested with 200 μL Tris-EDTA buffer (0.1 M Tris–HCl, pH 7.75 and 1 mM EDTA) and heated at 95 °C for 5 minutes. For the determination of ATP, 25 μL samples were mixed with 25 μL PEP buffer (5 mM PEP, 9 mM MgCl_2_, 5 mM KCl in 0.1 M Tris-EDTA buffer, PH 7.75).

### Mitochondrial membrane potential assay

As described previously (Zhang et al., 2018), mitochondrial membrane potential (Δψ_m_) was determined by flow cytometry-modulated JC-1 (5,5’,6,6’-tetrachloro-1,1’,3,3’-tetraethyl-benzimidazolylcarbocyanine iodide) assay according to the manufacturer’s instruction. Cells were harvested by 0.25% trypsin-EDTA, and incubated in cell media containing 5 μg/mL JC-1 (Enzo Life Sciences, NY, USA) for 15 min at 37 ^o^C. After washed once with PBS, samples were analyzed by a flow cytometer (FACSAria II, BD Biosciences, NJ, USA) using the excitation wavelength of 488 nm and the emission wavelengths of 525 nm for green fluorescence, or the emission wavelengths of 575 nm for orange-red fluorescence. The Δψ_m_ of each cell was calculated by the ratio of red fluorescence intensity to green fluorescence intensity. For each sample, the ratio of the cells with low Δψ_m_ in total cell population was reported by the flow cytometer (FACSAria II, BD Biosciences, NJ, USA).

### Statistical analyses

All data are presented as mean + SEM. Data were assessed by one-way ANOVA, followed by Student - Newman - Keuls post hoc test. *p* values less than 0.05 were considered statistically significant.

## Results

### Koisio media led to remarkable increases in the intracellular ATP levels of L929 cell cultures

We found that for the lung fibroblast L929 cell cultures, Koisio media led to approximately an 57% increase in the intracellular ATP levels of the L929 cell cultures of the Cell Passage 6 - 18, compared with regular cell culture media (Fig. 1).

**Fig. 1.**
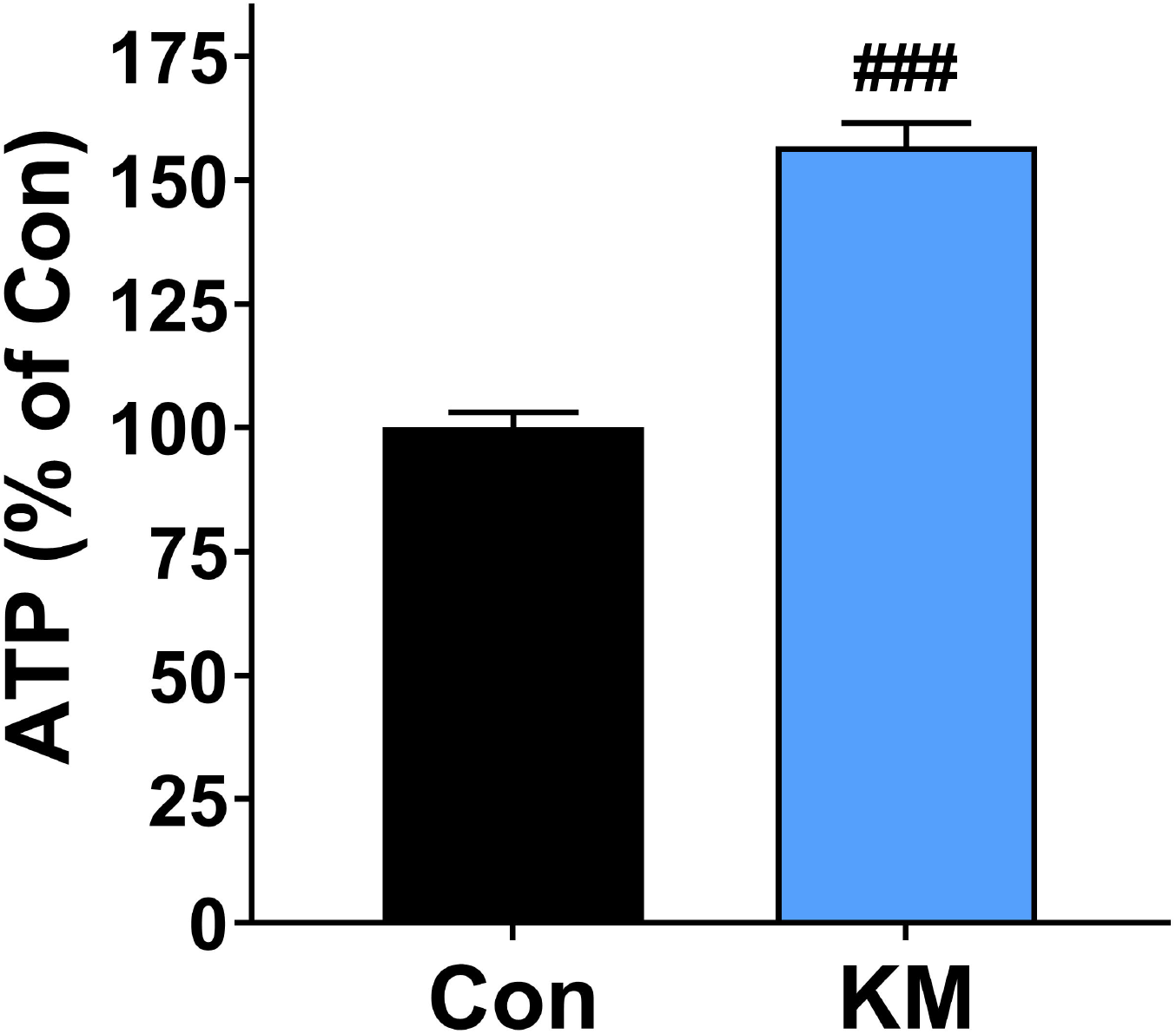
Koisio media (KM) led to remarkable increases in the intracellular ATP levels of L929 cell cultures. Cultures of the cells in Koisio media led to an approximately 57% increase in the intracellular ATP levels of the L929 cell cultures of the Cell Passage 6 – 18, compared with the cells cultured in regular cell culture media. The ATP assays were conducted 24 - 36 hrs after the cells were plated. N = 34. ###, *P* < 0.001 (Unpaired t-test).

**Fig. 2.**
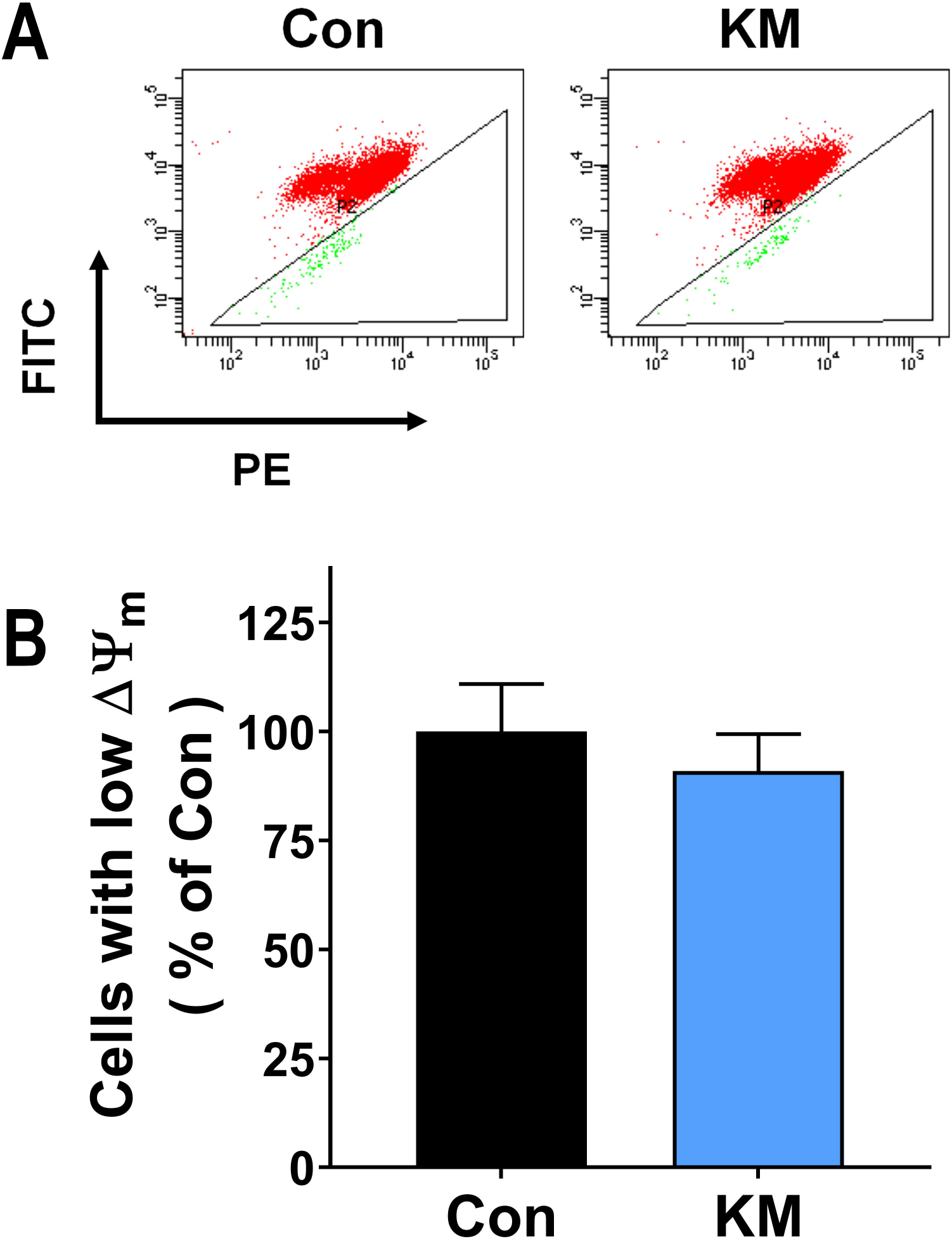
Koisio media (KM) did not significantly affect the mitochondrial membrane potential of the L929 cell cultures. Koisio media did not significantly affect the mitochondrial membrane potential of the L929 cell cultures of the Cell Passage 6 – 18, compared with the cell cultures cultured in regular media. The JC-1 assays were conducted 24 - 36 hrs after the cells were plated. N = 7.

### Koisio media did not significantly affect the mitochondrial membrane potential of L929 cell cultures

Twenty-four hr after the cells were cultured in Koisio media or regular cell cultures, JC-1 assays were conducted. We did not find that Koisio media could significantly affect the mitochondrial membrane potential of L929 cell cultures, compared with regular cell culture media (Fig. 2).

We also conducted experiments using another protocol of cell treatment: Twenty-four hr after the cells were cultured in Koisio media or regular cell culture media, the cells were cultured in MEM for 1 hr. Subsequently, JC-1 assays were conducted. In the study using this protocol, Koisio media did not significantly affect the mitochondrial membrane potential of L929 cell cultures (Fig. 3).

**Fig. 3.**
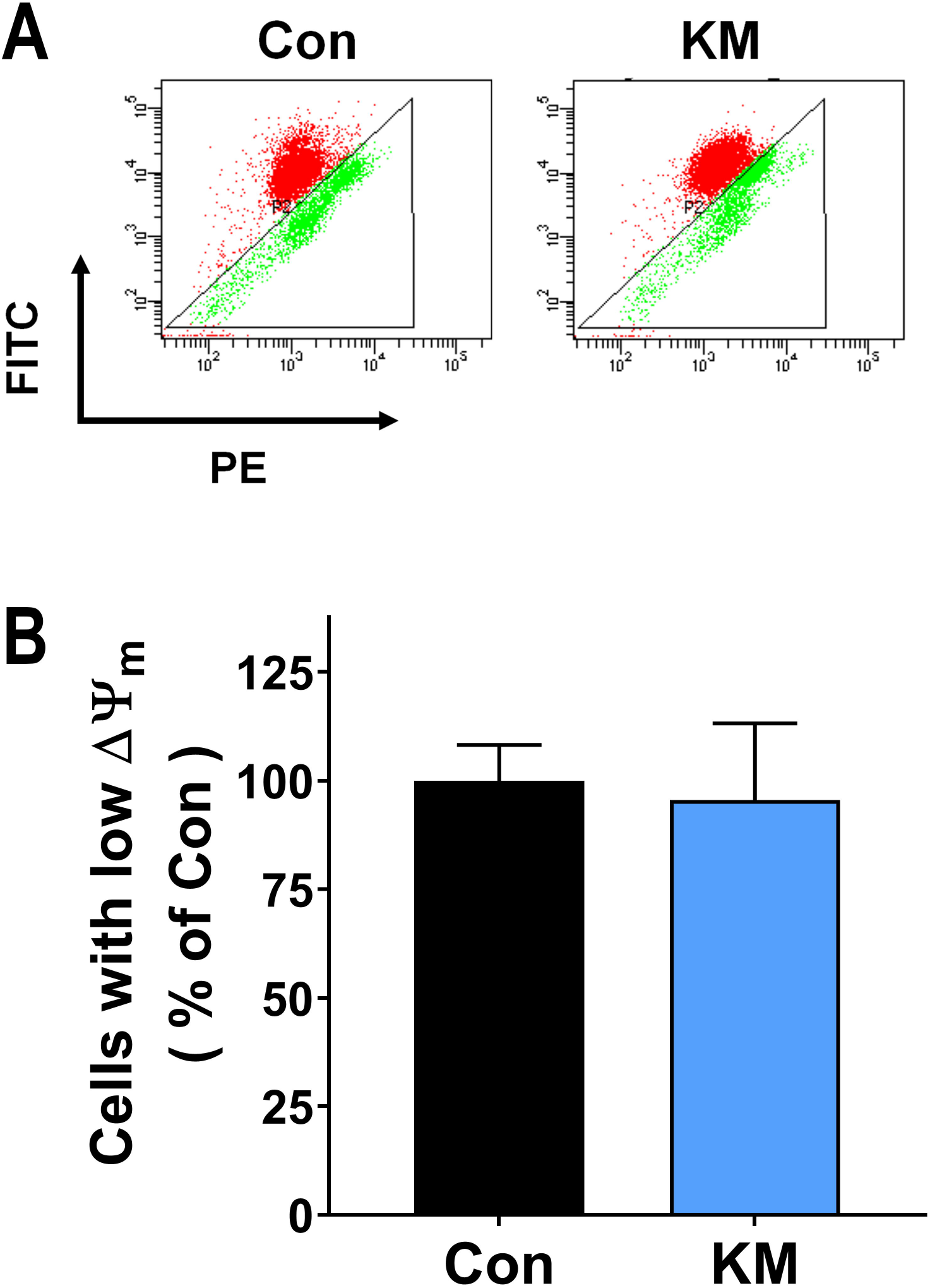
Koisio media (KM) did not significantly affect the mitochondrial membrane potential of the L929 cell cultures in a different cell culture protocol. Koisio media did not significantly affect the mitochondrial membrane potential of L929 cell cultures of the Cell Passage 6 – 18, compared with the cell cultures cultured in regular cell culture media. Twenty-four hr after the cells were cultured in Koisio media or regular cell culture media, the cells were cultured in MEM for 1 hr. Subsequently, JC-1 assays were conducted. N = 6.

## Discussion

The major findings of our study include: First, Koisio media led to approximately an 57% increase in the intracellular ATP levels of the L929 cell cultures of the Cell Passage 6 - 18, compared with regular cell culture media; and second, Koisio media did not significantly affect the mitochondrial membrane potential of the cell cultures, compared with regular cell culture media.

It is highly valuable to discover novel and economic substances for enhancing energy levels of human body. Since water drinking is required for human survival, it would be of great value if we can produce certain type of water with capacity of enhancing energy levels without addition of external substances. Our findings have shown that Koisio technology-modulated solutions have significant capacity to increase intracellular ATP levels. This finding has suggested that Koisio technology-modulated water could have significant capacity to increase intracellular ATP levels.

It is noteworthy that Koisio technology modulates water or solutions solely by physical modulations without additions of any external substances. To our knowledge, our finding that Koisio technology-modulated solutions have significant capacity to increase intracellular ATP levels is the first report that solutions modulated solely by physical approaches can have the capacity of increasing intracellular ATP levels. Future studies are needed to investigate the mechanisms underlying this novel finding.

Our study has provided first evidence indicating that Koisio media led to remarkable increases in the intracellular ATP levels of the L929 cell cultures without affecting mitochondrial membrane potential. This observation is seemingly surprising, considering the fact that mitochondrial oxidative phosphorylation is causative to the cellular generation of a large majority of ATP. These findings have suggested that Koisio media can lead to remarkable increases in intracellular ATP levels by novel energy-enhancing mechanisms. It is warranted to conduct future studies to investigate the novel mechanisms.

